# Adenovirus-based vaccines transduce and activate human Langerhans cells

**DOI:** 10.1101/2025.10.06.680090

**Authors:** Elina Gerber--Tichet, Thi Hong Giang Ngo, Fabien P. Blanchet, Eric J. Kremer

## Abstract

Langerhans cells (LCs) are specialized antigen-presenting cells (APCs) located in the epidermis and mucosal tissues and play a critical role in host immune surveillance. The role of LCs in antiviral responses is well established, but their involvement in vaccine-induced immunity, particularly with human adenovirus (HAdV) vectors, remains poorly understood. Here, we investigated how human LCs respond to HAdV vectors commonly used in vaccine development. We used CD34^+^ hematopoietic stem cells differentiated into LCs (CD34-LCs) to evaluate their ability to internalize replication-defective HAdV types C5, D26, and B35 vectors. CD34-LCs took up all three vectors, which induced an antiviral and pro-inflammatory cytokine response and morphological changes indicative of activation. To address the role of LCs in a more physiologically relevant setting, we injected human skin explants with the HAdV vectors. Epidermal LCs were recruited to the injection site and took up the vectors. Moreover, we show that the presence of lactoferrin (Lf), an antimicrobial protein, enhances HAdV uptake. Overall, our findings highlight a key role for LCs in the initiation of immune responses to HAdV-based vaccines. This work lays a foundation for strategies aiming to enhance vaccine efficacy by modulating LC activation and targeting.

**IMPORTANCE:** HAdV vectors are widely used in vaccine development, but the role of LCs, the epidermal immune sentinels, remains poorly defined. Here, we show using *in vitro* and *ex vivo* human systems that LCs efficiently detect, internalize and express transgenes delivered by three HAdV species. Because LCs link innate and adaptive immunity, the direct and indirect uptake of HAdVs may contribute to vaccine efficacy.

## INTRODUCTION

Over the last 50 years, vaccines have likely saved more than 154 million lives. Yet, we still poorly understand why some vaccines efficiently protect the host for decades (e.g., the tetanus/diphtheria/whooping cough vaccine), while others allow more modest short-term protection. An early component of vaccine efficacy is innate immune responses by antigen-presenting cells (APCs), especially those found in the skin. The skin provides a physical and immune barrier against external elements, including viral pathogens (1). Langerhans cells (LCs) are among the various innate immune cells that populate the skin (2, 3). In a healthy state, LCs are found exclusively in the epidermis and can be characterized their dendritic morphology, the presence of Birbeck granules and the expression of a C-type lectin receptor (CLRs) called langerin (CD207) (4, 5). CLRs are pattern recognition receptors (PRRs) which play key roles in pathogen recognition and immune response modulation. As sentinels, these APCs provide the link between innate and adaptive immunity (6–8) through the antigenic presentation process (9).

A handful of studies have described the response of LCs to viruses, including HIV-1 (10, 11), herpes simplex virus (12), dengue virus (13), and Usutu virus (14). A better understanding of the role of LCs during vaccination with HAdV-based vaccines could help improve their efficacy. HAdVs are non-enveloped particles belonging to the genus Mastadenovirus (15), with a linear double-stranded DNA genome of approximately 35 ± ∼5 kb (16, 17). HAdVs are classified into seven species (A to G), and exhibit a range of tissue tropisms and disease-related symptoms (18). Because human and nonhuman AdV-based vectors are rapidly and inexpensively produced, their use for long-term gene therapy (19, 20), antitumor therapy (21), and vaccines (22–26) compete with most vector platforms. Of note, during COVID-19 vaccination campaign HAdV-based vaccines were administered worldwide and protected millions of individuals (27).

Among the more developed vaccine candidates are HAdV-C5,-D26, and-B35 (18, 28, 29). In previous studies, we showed that HAdV-challenged human APCs show signs of functional maturation including changes in their transcriptome, changes in morphology, reduction in phagocytosis, and release of pro-inflammatory cytokines. Moreover, we and others showed that antimicrobial peptides (AMPs) released following neutrophil recruitment (30, 31), bind HAdV-C5,-D26, and-B35 via alternative receptors such as Toll-like receptor 4 (TLR4) and DC-SIGN (CD209) (32–34). This bridging effect enhances uptake of HAdV by certain human APCs. Recently, we catalogued and discussed the spectrum of receptors expressed by APCs that may mediate these interactions (35).

In this study, we expanded our understanding of the ability of HAdV-based vaccines to transduce and activate human LCs using *in vitro* and *ex vivo* approaches. LCs generated from CD34^+^ hematopoietic stem cells (CD34-LCs) took up HAdV-C5,-D26, and-B35 vectors directly and indirectly via lactoferrin (Lf). Capsid uptake was followed by cytokine secretion and morphological changes indicative of APC activation. In a second step, we used a physiologically relevant experiment: human skin explants from healthy donors, to reproduce a natural context of cutaneous vaccination. Administration of HAdV-based vaccines induced LC recruitment to the injection site, followed by vector uptake. Together, these studies support a model in which LCs play an initiating role in the establishment of immune responses to HAdV-based vaccines administered via the cutaneous route.

## MATERIALS AND METHODS

### Cell lines and culture conditions

CD34-LCs: CD34⁺ hematopoietic progenitors were isolated from umbilical cord blood using the EasySep CD34 Positive Selection Kit (STEMCELL Technologies) after Ficoll separation. Cells were expanded in StemSpan SFEM II medium with CC100 cytokine cocktail for 7 days. Then, cell differentiation into CD34-LCs was performed in RPMI 1640 with 10% FCS, GM-CSF (100 ng/mL), TGF-β (2 ng/mL), and TNF (2.5 ng/mL) for 10 days.

### HAdV vectors

The replication-defective, E1-deleted, HAdV vectors included HAdV-C5-GFP (36, 37), HAdV-D26-GFP (38), and HAdV-B35-YFP (39). The vectors were propagated in 911 or 293T E4-pIX cells and purified by CsCl gradient ultracentrifugation. Vector infectivity was assessed by GFP/YFP expression in permissive cells via flow cytometry.

### HAdV uptake by CD34-LCs

Cells (2 × 10⁵ per condition) were infected with HAdV-C5-GFP, HAdV-D26-GFP, or HAdV-B35-YFP at either 2.5 × 10³ or 1 × 10⁴ pp/cell. After 48 h, cells were harvested using 50 mM EDTA and analyzed by flow cytometry. After 24 h of *in vitro* exposure to the vaccine vectors, the CD34-LCs could be harvested for RNA extraction to analyze cytokine transcripts, while their culture supernatants were collected for cytokine release measurements.

### HAdV–lactoferrin complexes and inhibition assays

To assess indirect uptake, HAdV vectors were preincubated with 40 μg/mL Lf (cat#1294, Sigma-Aldrich) for 30 min at room temperature. HAdV-Lf complexes were added to CD34-LCs at 2 × 10⁴ pp/cell for 24 h. Inhibition of uptake was tested using 1 mg/mL mannan or intravenous immunoglobulin (IVIg; Baxter SAS).

### Flow cytometry

Cells were fixed in 2% paraformaldehyde and permeabilized with PBS containing 1% BSA and 0.05% saponin. Cells were stained with fluorophore-conjugated antibodies: CD207-PE (cat#130-098-355, Miltenyi biotec), CD1a-PECy5 (cat#300108, Biolegend), HLA-DR-APC (cat#130-113-398, Miltenyi biotec), and CD34-APC (cat#130-113-176, Miltenyi biotec), and acquired on a NovoCyte 3000RYB flow cytometer (Agilent). Data were analyzed with NovoExpress software.

### Surface plasmon resonance (SPR)

SPR was performed using a Biacore T200 instrument (GE Healthcare). HAdV capsids were immobilized on CM5S chips via amine coupling in 10 mM sodium acetate (pH 4.0). Immobilization levels ranged from 985 to 5192 RU. To determine the impact of Lf, recombinant human langerin protein (cat#2088-LN, R&Dsystem) (hCD207) and Lf (cat#1294, Sigma-Aldrich) were injected at 100 nM. To determine the K_D_ between HAdVs and langerin, different concentrations of langerin (6.25–200 nM) were injected for 120 s (association) and 400 s (dissociation) at 30 μL/min. The kinetic constants were evaluated from the sensorgrams after subtraction of flow cell 1 using BIAevaluation 3.2 software (GE Healthcare).

### Quantification of *IL1B, IFNB, CXCL10* and *CCL5* mRNAs

Total RNA was extracted using the RNeasy Mini Kit (Qiagen), and cDNA was synthesized using the SuperScript VILO cDNA Synthesis Kit (cat#11754-050, Thermo Fisher Scientific). Quantitative PCR was performed with PowerUp SYBR Green Master Mix (cat#A25742, Thermo Fisher Scientific) using a LightCycler 480 (Roche). Expression of *IFNβ*, *CCL5*, *CXCL10*, and *IL-1β* was normalized to *GAPDH* using the 2^–ΔΔCt method.

### Cytokine secretion

Cytokine levels (CXCL10, CCL5, IFNβ, and IL-1β) in culture supernatants were quantified using ELISA MAX Deluxe Kits (BioLegend) per the manufacturer’s instructions.

### Scanning electron microscopy (SEM)

CD34-LCs were cultured on 0.1% poly-L-lysine–coated coverslips (4 × 10⁵ cells/500 μl of complete medium). Cells were incubated with 2 × 10⁴ pp/cell of viral particles for 24 h, then fixed with 2.5% glutaraldehyde in 1× PHEM buffer (240 mM PIPES, 100 mM HEPES, 8 mM MgCl₂ ·6H₂O, 40 mM EGTA). Samples were dehydrated in a series of increasing ethanol concentrations (30, 50, and 70%). Sample preparation for SEM was performed by the COMET electron microscopy core facility of the Institute of Neurosciences of Montpellier (INM, University of Montpellier/INSERM), and images were acquired using a Hitachi S4000 scanning electron microscope.

### Atomic force microscopy (AFM)

CD34-LCs were seeded on 0.1% poly-L-lysine–coated coverslips (1 × 10⁵ cells/500 μl complete medium) and exposed to 2 × 10⁴ pp/cell for 24 h. Topographical imaging and mechanical characterization were conducted using a Nanowizard 4 AFM (JPK-Bruker). A tip mounted on a flexible cantilever was used to apply point-by-point force on the cell surface. Deflection of the laser beam reflected off the cantilever was measured by an optical detector to map the three-dimensional (x, y, z) surface topography. The force-distance curves obtained were used to calculate the Young’s modulus (YM), which represents cellular elasticity.

### Human skin processing

Human abdominal skin biopsies were collected from patients receiving abdominoplasty at the Montpellier University Hospital, France, with informed consent. Experiments were conducted under agreement [DRI_2021-58] – BDT-Peaux-Dr BLANCHET / CNRS Ref. 249300 and with support from the CRB-CHUM biobank platform (http://www.chu-montpellier.fr; Biobank ID: BB-0033-00031). Subcutaneous fat was removed, and 1 cm² skin explants were cultured at an air-liquid interface in RPMI-1640 medium supplemented with 10% FCS.

### Vector injection in human skin explants

Human skin explants were injected intradermally using 29G insulin syringes with 1 × 10¹¹ pp of human HAdV vectors (HAdV-C5,-D26, and-B35). After 48 h, tissues were fixed in 10% paraformaldehyde for 24 h at 4°C, dehydrated in 70% ethanol, and paraffin-embedded by the experimental histology platform (RHEM) at the Institut de Recherche en Cancérologie de Montpellier (IRCM).

### Immunohistochemistry (IHC) on paraffin-embedded skin

Ten-micrometer sections were cut using a Microm HM340E microtome equipped with a Niagara system at the RHEM platform of the IGMM. Sections were deparaffinized and rehydrated through xylene and ethanol washes, followed by antigen retrieval in 10 mM citrate buffer at 100°C for 10 min. Endogenous peroxidase activity was blocked with H₂O₂ for 20 min. Tissue sections were pre-incubated for 30 min in antibody diluent to reduce background and increase permeability, before being incubated overnight at 4°C with primary anti-CD207 antibody (cat# HPA011216, Sigma; 1:150 dilution). For immunohistological detection, sections were incubated for 1 h with a biotinylated secondary antibody at room temperature in the dark, followed by avidin-biotin-peroxidase complex and DAB substrate development. Slides were mounted using Dako mounting medium (cat#S3023) and imaged using a Leica Thunder upright microscope at the Montpellier Ressources Imagerie (MRI) core facility.

### Immunofluorescence (IF) on paraffin-embedded skin

Following the same protocol as IHC, primary antibodies used included anti-CD207 (cat# HPA011216, Sigma), anti-CD5 (cat# AMAb91782, Sigma), and anti-CD206 (cat# AF2534, R&D Systems), each diluted 1:150. Secondary antibodies conjugated to fluorophores were used at a 1:200 dilution. Nuclei were stained with DAPI. Slides were mounted with Dako medium (cat# S3023) and visualized with a Leica Thunder upright or a Zeiss LSM980 confocal microscope at the MRI platform (CRBM). Images were analyzed using QuPath software to quantify cell density (cells/cm²) in regions surrounding the injection site.

## RESULTS

### CD34-derived human Langerhans cells (CD34-LCs)

To generate an *in vitro* model of human LCs, CD34⁺ hematopoietic stem/progenitor cells (HSCs) were isolated from umbilical cord blood mononuclear cells and subsequently expanded. The HSCs were then incubated with GM-CSF, TGF-β, and TNF for 10 days to induce their differentiation into LCs. Under these conditions, the cells expressed hallmark surface markers of LCs and acquired typical dendritic morphology. They also formed cell aggregates through E-cadherin–mediated intercellular adhesion, due to intercellular adhesion mediated by E-cadherin in response to TGF-β signaling (**Figure** 1A) (40, 41). Undifferentiated HSCs were CD1a^-^, CD207^-^, and HLA-DR^low^ (42).

**Figure 1.**
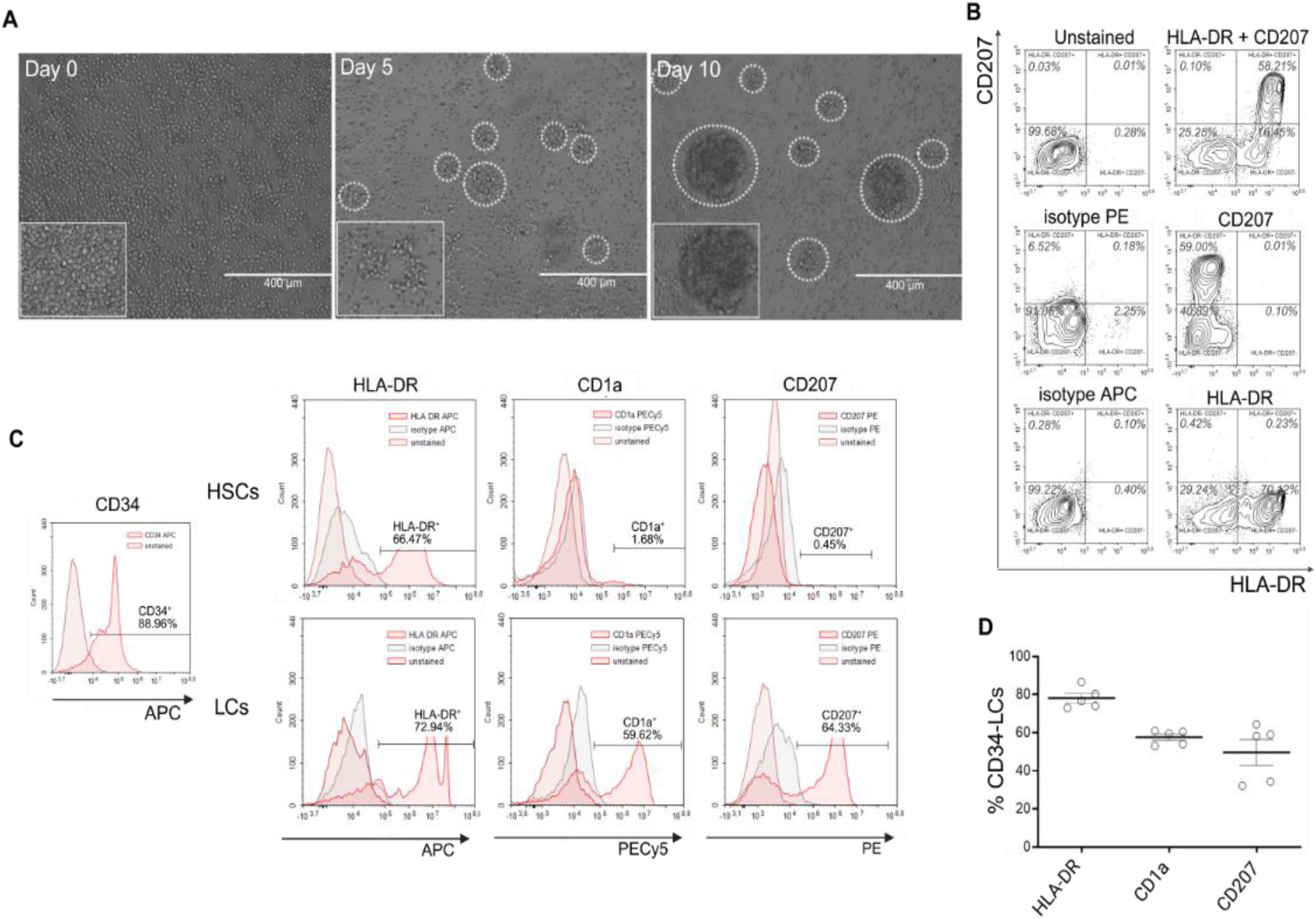
Differentiation strategy, phenotypic characterization, and validation of CD34-LCs. (A) Brightfield microscopy images of cells undergoing differentiation at day 0 (beginning), day 5 (half differentiation) and day 10 (end differentiation). (B) Flow cytometric analysis of the co-expression of HLA-DR and CD207 in CD34-LCs. The presence of these markers confirms the antigen presentation function coupled with LC phenotype obtained through 10 days of cell differentiation. (C) Flow cytometric phenotyping of CD34-LCs and their precursors, showing the expression of each specific identification markers (HLA-DR, CD1a, CD207). (D) Percentage of CD34-LCs expressing HLA-DR, CD1a, or CD207 in the final differentiated cell population in all donors, as determined by flow cytometry.

All CD207⁺ co-expressed HLA-DR, while some HLA-DR⁺ cells remained CD207^-^, showing that langerin is restricted to cells with an APC phenotype. The presence of HLA-DR⁺/CD207⁻ cells likely reflect incomplete or asynchronous differentiation, a phenomenon expected in stem cell-derived systems (**Figure** 1B). Flow cytometry analysis revealed that >70% of cells expressed HLA-DR, and around 60% co-expressed CD1a and CD207, consistent with the generation of an LC-like population (**Figure** 1C) (43, 44). CD34-LCs from several donors had similar profiles (**Figure** 1D). Finally, comparative phenotypic analysis revealed that CD34-LCs exhibited significantly higher CD207 expression than other LC-like models, including primary epidermal LCs (eLCs), monocyte-derived LCs (moLCs) and CHO cells transfected with CD207 (CHO-207) (**Figure** S1A). Both, the frequency of CD207⁺ cells and the fluorescence intensity of CD207 staining, were markedly higher in CD34-LCs.

### HAdV uptake and transgene expression by CD34-LCs

CD34-LCs were incubated with 2.5 × 10 or 1 × 10⁴ ³ physical particles (pp)/cell of HAdV-C5,-D26, and-B35 for 48 h based on preliminary analyses of optimal infectivity for this cellular model (data not shown). GFP or YFP expression, detected by flow cytometry, was used as a surrogate for HAdV uptake (**Figure** 2).

**Figure 2.**
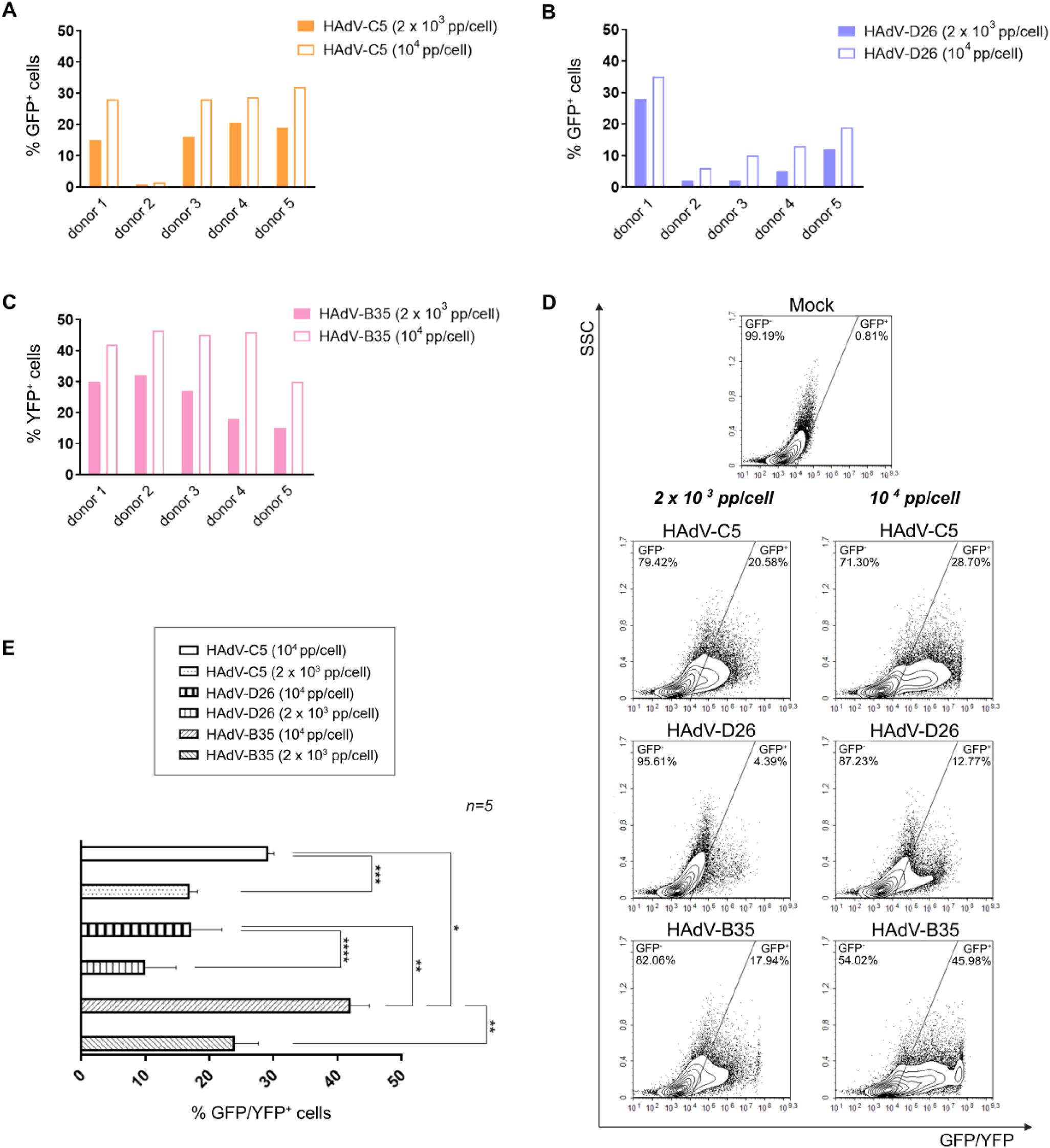
CD34-LCs take up HAdV-C5,-D26, and-B35. (A–C) Flow cytometry analysis of CD34-LCs taking up HAdV-C5 (A), HAdV-D26 (B), or HAdV-B35 (C) (n = 5 donors) 48 h post-exposure. Two doses (2.5 × 10³ pp/cell or 1 × 10⁴ pp/cell) were used, and uptake was assessed by GFP or YFP transgene expression. (D) Representative flow cytometry profiles showing GFP/YFP expression in CD34-LCs after 48 h of exposure to HAdV vaccine vectors at the indicated doses. (E) Comparison of HAdV-C5, - D26, and-B35 uptake across all donors. (*p < 0.05 to ****p < 0.0001)

Although there was inter-donor variability, a trend of transduction efficiency was similar in all donors with HAdV-C5 (**Figure** 2A),-D26 (**Figure** 2B) and-B35 (**Figure** 2C). CD34-LCs internalized the vectors with different efficiencies (**Figure** 2D): At 2.5 × 10³ pp/cell, no major differences were observed between vectors, while at 10⁴ pp/cell, HAdV-B35 showed the highest transduction efficacy (mean: 42% YFP⁺), followed by HAdV-C5 (30% GFP⁺), and HAdV-D26 (18% GFP⁺) (**Figure** 2E). These results showed that CD34-LCs are capable of internalizing and expressing transgenes from HAdV-C5,-D26 and –B35 replication-defective vectors.

### Pro-inflammatory and antiviral response of CD34-LCs

To characterize the innate immune response of CD34-LCs following exposure to HAdV-C5,-D26,-B35, we quantified the mRNA expression from genes encoding pro-inflammatory and antiviral cytokines. Twenty-four hours post-challenge with 2.5 × 10³ or 1 × 10⁴ pp/cell we observed an increase of *IL1B, IFNB1, CXCL10,* and *CCL5* transcripts (**Figure** 3A).

**Figure 3.**
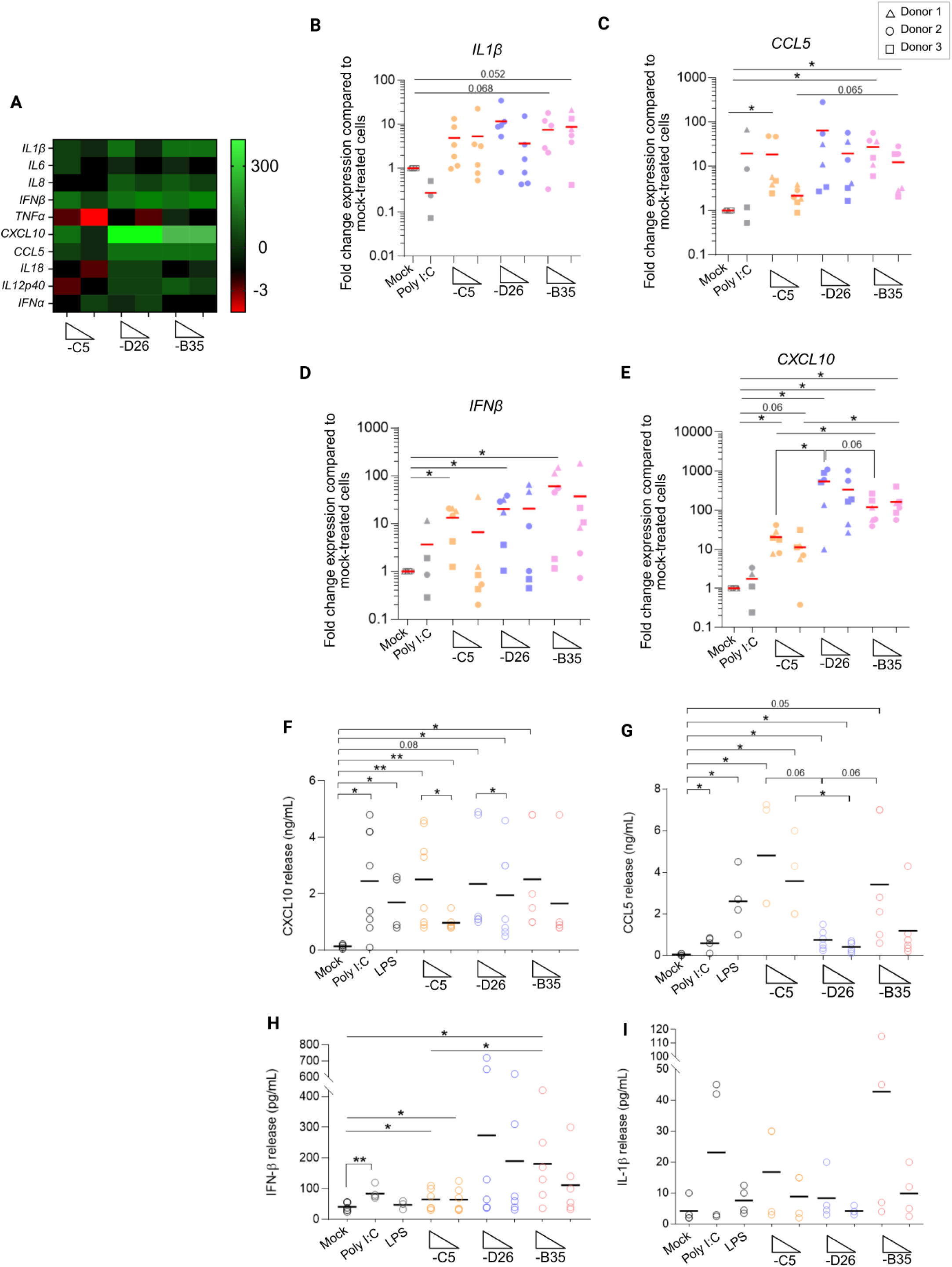
HAdV vectors induce a pro-inflammatory and antiviral state in CD34-LCs. (A) Heatmap showing changes in transcript levels of pro-inflammatory and antiviral cytokines in CD34-LCs after exposure to HAdV-C5, HAdV-D26, or HAdV-B35 at the indicated doses, compared to the mock-treated cells. Red shading indicates unchanged transcript levels, whereas green shading denotes increased transcript levels in response to vector exposure. (B-E) Quantitative analysis by RT-PCR of pro-inflammatory and antiviral cytokine transcripts (CXCL10 (B), CCL5 (C), IFN-β (D), and IL-1β (E)) in CD34-LCs after exposure to HAdV-C5, - D26, and-B35 vectors for 24 h at doses 2.5 × 10³ pp/cell or 1 × 10⁴ pp/cell. The expression of IFNβ, CCL5, CXCL10, and IL-1β was normalized to GAPDH using the 2^–ΔΔCt method. (F-I) Level of secretion of CXCL10 (F), CCL5 (G), IFN-β (H) and IL-1β (I) proteins by CD34-LCs that had been in contact with HAdV-C5,-D26, and-B35 at doses 2.5 × 10³ pp/cell or 1 × 10⁴ pp/cell for 24 h, assessed by ELISA. (*p < 0.05 to ****p < 0.0001)

Of note, these cytokines are also upregulated in human moDCs challenged with HAdV-C5,-26 and-B35 vectors (32, 33, 45–48). Exposure to all three vectors appears to induce a tendency to increase *IL1B* mRNA at both doses, with a change closer to significance for HAdV-B35 (**Figure** 3B). The changes in *CCL5* mRNA levels (**Figure** 3C) were significant for HAdV-C5 at 1 × 10⁴ pp/cell (∼ 20 fold) and also-B35 at both doses tested (between 20 and 30 fold).

*IFNB1* mRNA levels increased between 7 and 50 fold, with significant induction by all three HAdVs at 1 × 10⁴ pp/cell only. This demonstrated a possible dose-dependent increase in *IFNB* transcript levels (**Figure** 3D). Similarly, the presence of each vector at 1 × 10⁴ pp/cell induced an increase in *CXCL10* mRNA. However, unlike the other cytokine mRNAs studied, it appears that *CXCL10* mRNA showed vector-type-dependent variations. Thus, at 1 × 10⁴ pp/cell, HAdV-D26 induced the highest increase (approximately 600 times higher) and-C5 the lowest (approximately 10 to 20 times higher) (**Figure** 3E).

ELISA analysis of supernatants confirmed the production of the corresponding cytokines, even if the results for transcript levels and cytokine production quantities were not comparable.

All three vectors led to a significant secretion of CXCL10 (1-3 ng/mL), in a dose-dependent manner, except for HAdV-B35 (**Figure** 3F).

HAdV-C5 induced the highest CCL5 production (between 3.5 and 5 ng/mL), with concentrations 30 to 50 times higher than in mock-treated cells; and-D26, the lowest CCL5 production (<1ng/mL). But overall, HAdV-C5,-D26 and –B35 induced a significant increase in CCL5 (**Figure** 3G). At the protein level, although all vectors showed an overall tendency toward increased IFN-β secretion, only certain conditions were significantly different: HAdV-C5 at 2.5 × 10³ and 1 × 10⁴ pp/cell (∼50 pg/mL), and HAdV-B35 at 1 × 10⁴ pp/cell (100 to 200 pg/mL) (**Figure** 3H). Thus, despite some differences in significance at the protein level, our data suggest that all three vectors induce a general trend toward increased IFN-β.

With regard to IL-1β, although concentrations were in low concentration ranges like IFN-β, HAdV-B35 induced secretion up to more than 40 pg/mL (i.e. ∼10 times greater than the control condition). But none of these changes revealed really significant differences compared to mock-treated cells, as also observed in *IL1B* mRNA levels (**Figure 3I**).

As a result, it would appear that IL-1β, produced by the inflammasome, is the least secreted or transcribed. Here, we demonstrate that CD34-LCs are able to detect HAdV-C5,-D26 and-B35 vectors and initiate a robust pro-inflammatory and antiviral response through changes in transcriptional and protein levels. These suggest that LCs initiate the local immune response following cutaneous administration of HAdV-based vaccines, thereby contributing to vaccine efficacy.

### HAdV-induced CD34-LCs activation correlates with morphological changes

In addition to transcriptional and cytokine changes, cellular morphology is an indicator of APC activation. Upon maturation, LCs undergo cytoskeletal reorganization that reduces antigen uptake and facilitates their migration to lymph nodes for antigen presentation. To investigate whether HAdVs trigger such morphological changes, we examined CD34-LCs by scanning electron microscopy (SEM) 24 h post-challenge with HAdV-C5,-D26, or-B35 (**Figure 4A**). Mock-treated CD34-LCs possessed multiple dendrites, which are characteristic of immature LCs capable of sampling their environment. By contrast, HAdV-stimulated CD34-LCs showed a marked retraction of dendrites and adopted a rounded, smooth morphology, consistent with functional maturation. This phenotype was also observed upon stimulation with low-and high-molecular-weight poly I:C, but not, as expected with LPS (LCs do not express TLR4) (**Figure 4A**). It should be noted that a similar morphological change has been described after contact of moDCs with HAdV-C5 (47).

**Figure 4.**
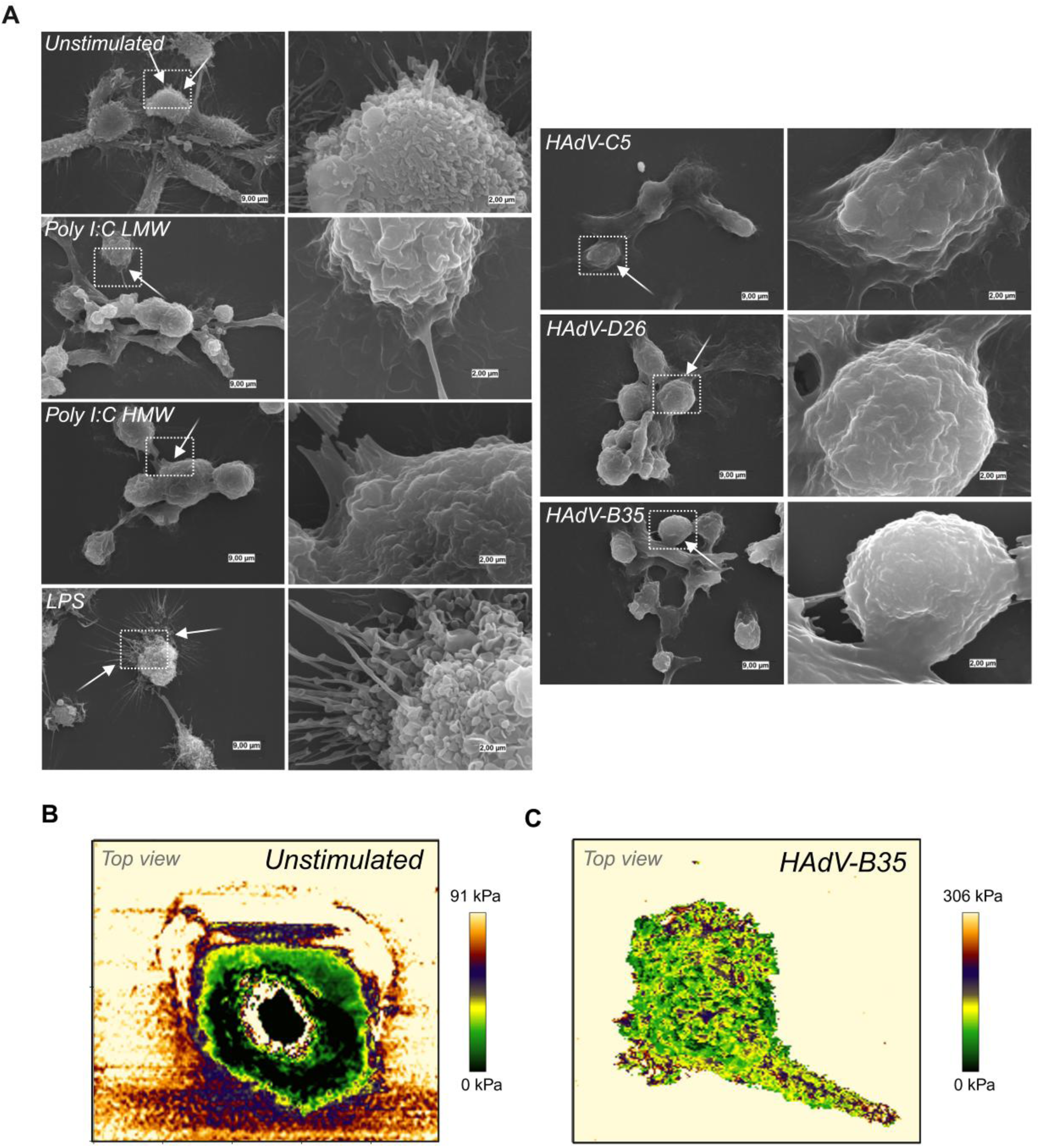
HAdVs increase cell rigidity and reduced dendrites in CD34-LCs. (A) SEM showing morphological changes in CD34-LCs following 24 h incubation with 2 × 10⁴ pp/cell of HAdV-C5,-D26,-B35, poly I:C LMW and HMW, or LPS (n = 3). Areas denoted by dotted lines are magnified in the right panel (B) AFM images of unstimulated (B) and HAdV-B35-stimulated cells (C); elasticity (kilopascal, kPa) is color-coded from black (more elastic) to beige (less elastic).

To evaluate whether this morphological remodeling corresponded to biophysical changes in cell rigidity, we performed atomic force microscopy (AFM) on unstimulated CD34-LCs or challenged with HAdV-B35 to measure cell stiffness (**Figure 4B-C**). While unstimulated cells had an average Young’s modulus (YM) of ∼120 kPa, HAdV-stimulated CD34-LCs had a higher Young’s modulus (∼ 250 kPa). In this representative image, the deformation forces ranged from 0 to 306 kPa in the case of HAdV-B35 stimulation (**Figure 4C**), indicating increased mechanical rigidity (49).

### Lf-and IVIg**-**enhanced HAdV uptake by CD34-LCs

Lf is a multifunctional and iron-binding glycoprotein with antiviral and immunomodulatory properties. We previously demonstrated that Lf can act as a bridging factor between moDCs and HAdVs to increase their uptake via toll-like receptor 4 (32, 33, 50). To determine if CD34-LCs uptake of HAdVs is also enhanced by Lf, we incubated them with HAdV-C5,-D26 or-B35 vectors alone, or complexed with Lf and/or IVIg. IVIg is made up of IgGs from several thousand North Americans donors and contains a significant level of HAdV-C5 neutralizing antibodies (NAbs), a moderate to low level of HAdV-D26 NAbs, and very low or no HAdV-B35 NAbs (18, 51). Using flow cytometry and transgene expression (GFP or YFP) expression as a readout, we found that Lf enhanced the uptake of all three HAdVs by CD34⁺ LCs (**Figure 5A-B**) with a particularly strong effect observed for HAdV-B35. As expected, pre-incubation of HAdV-C5 with IVIg led to a strong reduction of its uptake by CD34-LCs, while only a minor decrease was observed for HAdV-D26. In contrast, IVIg had no effect on the uptake of HAdV-B35, consistent with the lack of HadV-B35 NAbs (18, 52–54). When Lf and IVIg were co-incubated with the HAdVs, uptake of HAdV-C5 and HAdV-D26 was partially or totally restored, respectively (**Figure 5A-B**). Nevertheless, some uptake remained, as Langerhans cells (LCs) express Fcγ CD64 (FcγRI), CD32 (FcγRII), and CD16 (FcγRIII) receptors (52), known to be involved in the internalization of immune complexes (ICs) via the Fc region of IgG (53). These results highlight the potential of Lf to influence HAdV recognition by LCs.

**Figure 5.**
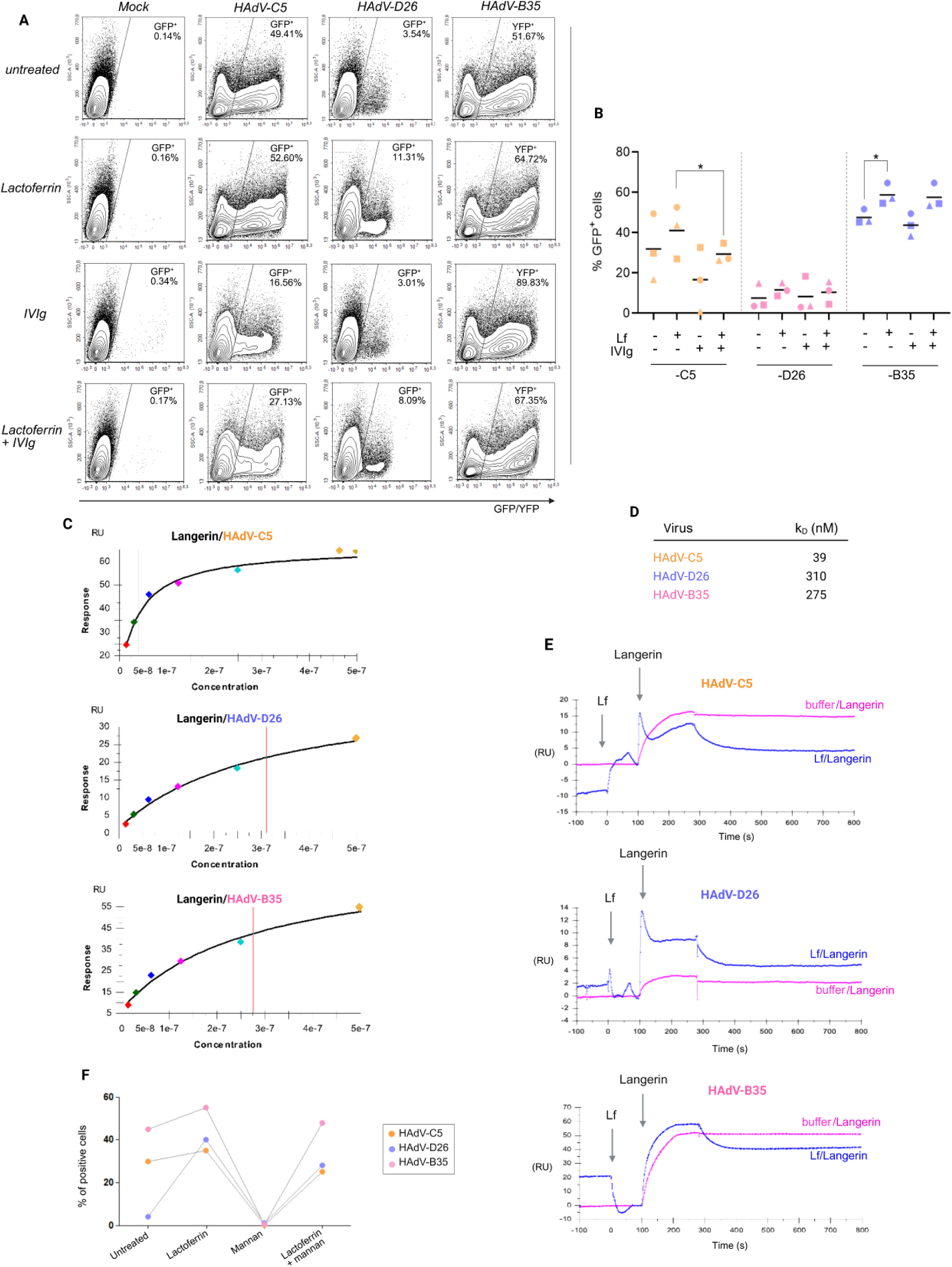
Lf enhances HAdVs uptake by CD34-LCs, independent of langerin. (A) Flow cytometry analyses of CD34-LCs uptake of HAdV-C5,-D26, and-B35 ± Lf and/or IVIg. HAdV-C5,-D26, and-B35 are replication defective vectors harbouring a fluorescent protein expression cassette. (B) Cumulative data from four CD34-LC donors. (C) Surface plasmon resonance (SPR) analysis of direct binding between langerin and immobilized HAdV-C5,-D26, and-B35 vectors. (D) Dissociation constants (K_D_) indicating the relative affinity of langerin for each vector. (E) Effect of Lf on langerin–virus interactions measured by SPR. (F) Impact of langerin blockade on vector uptake, showing that Lf-enhanced infection occurs independently of langerin engagement. (*p < 0.05 to ****p < 0.0001)

### Langerin - an additional receptor for HAdV-C5 and-D26?

As highly phagocytic cells that regularly sample the extracellular environment, LCs likely take up pathogens, including HAdVs, by receptor-independent processes. Moreover, while HAdV-B35 primarily uses CD46 as a receptor, HAdV-C5 and-D26 can bind multiple cell surface proteins to initiate uptake (35, 55). An additional layer of complexity is the role of AMPs, like Lf, which retarget HAdVs to alternative cell surface proteins such as TLR4 and DC-SIGN (CD209). Unlike dermal dendritic cells (dDCs), moDCs and moLCs, CD34-LCs have an undetectable level of TLR4 on their surface (56, 57). Langerin (CD207), like DC-SIGN, belongs to the C-type lectin receptor (CLR) family and is specifically expressed by LCs. We therefore asked if langerin could directly, or indirectly via a Lf bridge, promote HAdVs uptake. Using surface plasmon resonance (SPR), we found that langerin could directly interact with HAdV-C5,-D26, and-B35 capsids in a dose-dependent manner (**Figure 5C**), but only a small amount of molecules were bound to the surface of the three vectors. In contrast, a K_D_ of 39 nM, close to the binding range of CD46 to the-B35 fiber knob (15 nM) (58), was indicative of a strong affinity between langerin and HAdV-C5 (**Figure** 5D). O-GlcNAc modifications in the HAdV-C5 fiber shaft could be recognized by lectin-like receptors (59–61) and could explain the high-affinity binding specific to HAdV-C5. However, it should be noted that we do not know whether HAdV-D26 or-B35 fibers also contain O-GlcNAc modifications.

Next, we asked if Lf can influence HAdV-langerin interactions. Using a dual-injection SPR approach, we bound HAdV capsids to the chip, then injected Lf followed by langerin. We observed no change in the binding between langerin and HAdVs, suggesting that the enhancing effect of Lf on vector uptake is not due to an increased interaction via langerin (**Figure** 5E). To address this using another approach, we use mannan to saturate CLRs like langerin on the surface of CD34-LCs.This treatment seemed to abolish the uptake of HAdVs, validating the essential role of CLRs in direct recognition. However, pre-complexing HAdVs with Lf restored vector internalization even in the presence of mannan, suggesting that Lf redirects HAdVs toward non-CLR receptors (**Figure** 5F). These observations showed two distinct new pathways for HAdV vector uptake by CD34-LCs: a direct route via various receptors including langerin, and an indirect route mediated by Lf.

### Lf-mediated HAdV uptake induces a comparable activation profile to direct entry in CD34-LCs without causing hyperactivation

To characterize the innate immune response triggered by the Lf complexes, we analyzed cytokine mRNA and protein levels following incubation with HAdV-C5,-D26, or-B35 ± Lf. It should be noted that the induction of the expression of certain cytokines or the increase in their transcripts with LPS can be explained by the fact that, although LPS is not the most effective inducer of inflammation in this cellular context, given that the TLR4 receptor is weakly expressed and its role is widely debated, it appears capable of partially activating certain signaling pathways.

*CXCL10* mRNA expression was induced 10 - 40 fold with the HAdVs alone and further seem to increase up to 60-fold when Lf complexes were used (**Figure** 6A). At the protein level, CXCL10 (IP-10) secretion increased in response to HAdV vectors, and this secretion was amplified when vectors were complexed with Lf, except for HAdV– B35 (**Figure** 6B).

**Figure 6.**
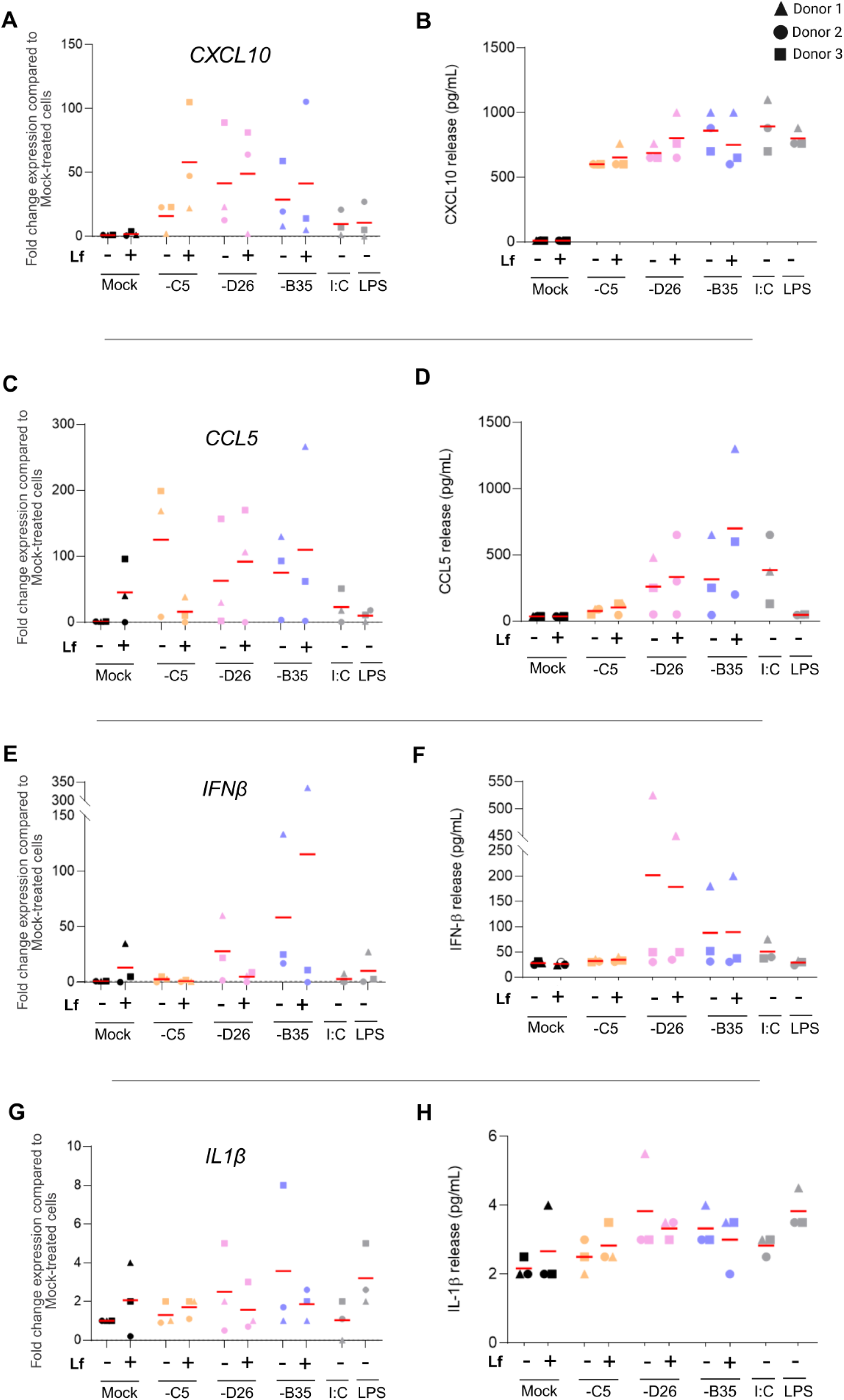
Effect of HAdV - Lf complexes on the activation of CD34-LCs. mRNA levels were measured by RT-qPCR and normalized to GAPDH using the 2^–ΔΔCt. The corresponding protein was quantified in culture supernatants by ELISA. The selected cytokines are pro-inflammatory and antiviral. Panels show mRNA expression and protein release for CXCL10 (A and B respectively), CCL5 (C and D respectively), IFN-β (E and F respectively) and IL-1β (G and H respectively) in CD34-LCs exposed to HAdV-C5, HAdV-D26, and HAdV-B35 during 24 h, either alone or pre-complexed for 30 min at room temperature with 40 μg/mL Lf. Experimental controls include mock-treated cells and cells treated with poly I:C (I:C) or lipopolysaccharide (LPS) as canonical innate immune stimuli. Each dot represents an individual donor; red lines indicate the mean for each condition. Data are representative of at least n = 3 independent donors. (*p < 0.05 to ****p < 0.0001)

Similarly, *CCL5* transcripts appeared to be upregulated by the HAdVs+Lf (**Figure** 6C). This transcriptional activation translated into increased CCL5 (RANTES) secretion, with a little enhancing effect observed in the presence of Lf (**Figure** 6D).

With Lf, IFN-β transcripts remained broadly unchanged for HAdV-C5, slightly decreased for HAdV-D26, and moderately increased for HAdV-B35 (**Figure** 6E). At the protein level, a similar profile was observed for HAdV-C5+Lf and-D26+Lf, while HAdV-B35+Lf did not induce any notable changes in IFN-β secretion, compared to HAdV alone (**Figure** 6F).

To finish, IL-1β transcription (**Figure** 6G) and secretion (**Figure** 6H) show an increasing trend for HAdV-C5 in the presence of Lf, whereas for HAdV-D26 and-B35, Lf tended to induce a decrease, as previously observed by our team with moDCs (32). These observations are consistent with the strong affinity of Lf for HAdV-C5, as previously described (32). It could promote the redirection of these viruses to receptors capable of activating the NF-κB and inflammasome pathways (e.g., TLRs, CLRs…) as observed with TLR4 on moDCs.

It should be noted that none of the observed results were statistically significant and instead represent trends. These results suggest that HAdV vectors complexed with Lf maintain the secretion of anti-viral chemokines, with a potential limited enhancing impact on inflammasome-dependent cytokines such as IL-1β (62). The data support an immunomodulatory role for Lf in the immune response to HAdVs, potentially through the selective engagement of certain receptors on the surface of CD34-LCs. However, even though Lf increases vector uptake by CD34-LCs, it does not appear to induce a stronger immune response in these cells under our conditions.

### LCs in human skin explants are mobilized and take up HAdVs

To analyze the impact of HAdVs on LCs using another approach, we injected the vectors into human skin explants obtained after abdominoplasties. After injection of HAdV-C5,-D26 or-B35 vectors, the explants were cultured for an additional 48 h, fixed, and then analyzed by immunofluorescence.

In uninjected skin, CD207⁺ LCs were distributed throughout the epidermis, mainly in the suprabasal layers, with no particular clustering (**Figure** 7A). By contrast, HAdV injections induced a relocalization of LCs, forming clusters at the injection site (**Figure** 7B). Taken together, these results highlight the ability of LCs to interact and uptake HAdV vectors in human skin *ex vivo*, suggesting their potential role as sentinel but also initiators of immune responses to HAdV-based vaccines, confirming the previous observations made *in vitro*.

**Figure 7.**
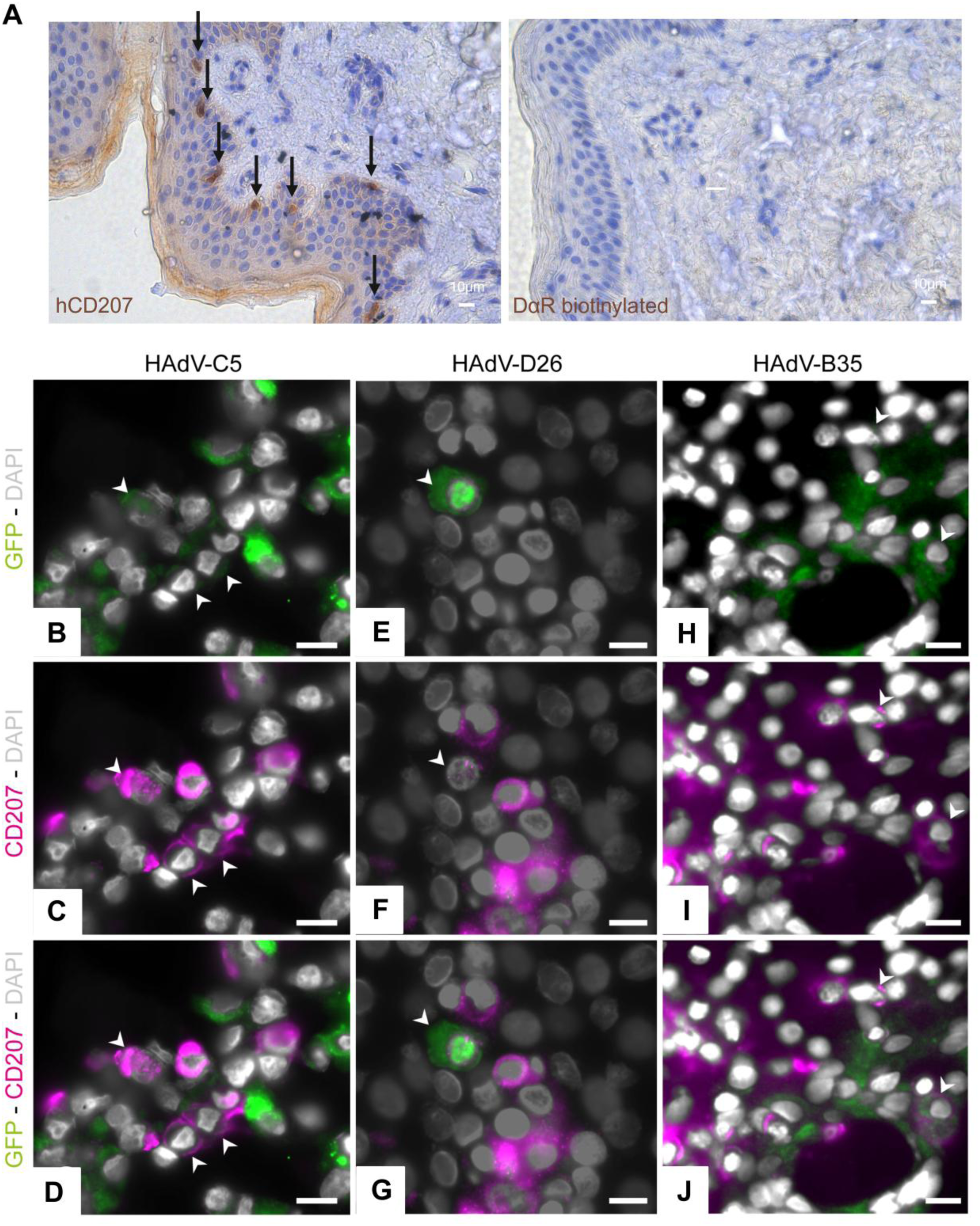
Primary epidermal LCs are transduced by HAdVs. (A) CD207⁺ immunoreactivity in healthy human skin explants. LCs are denoted by black arrows. The right-hand panel shows staining control using the secondary antibody. Representative images showing LCs located near the injection site capturing HAdV-C5 (B-D),-D26 (E-G), or-B35 (H-J), visualized by the colocalisation GFP/YFP reporter expression (green) and CD207 staining (magenta) (scale bar = 10 μm). White arrows show LCs that have internalized HadVs.

### HAdV injections promote the recruitment of LCs and CD5⁺ APCs

Firstly, we wanted to better characterize the identity and localization of antigen-presenting cells (APCs) in human skin. So, we performed multiple immunofluorescence staining for the markers CD207, CD5 and CD206 (**Figure** 8A). During the observation, LCs were confined to the epidermis, with an occasional presence in hair follicles, while cDC1s which are CD206⁺CD5^-^, were localized in the dermis and follicles, and absent from the epidermis. CD206⁺CD5⁺ cells, associated with migrating cDC2s, were mainly detected at follicular structures in basal condition, suggesting spatial segregation between LCs, cDC1s and cDC2s (**Figure** 8B).

**Figure 8.**
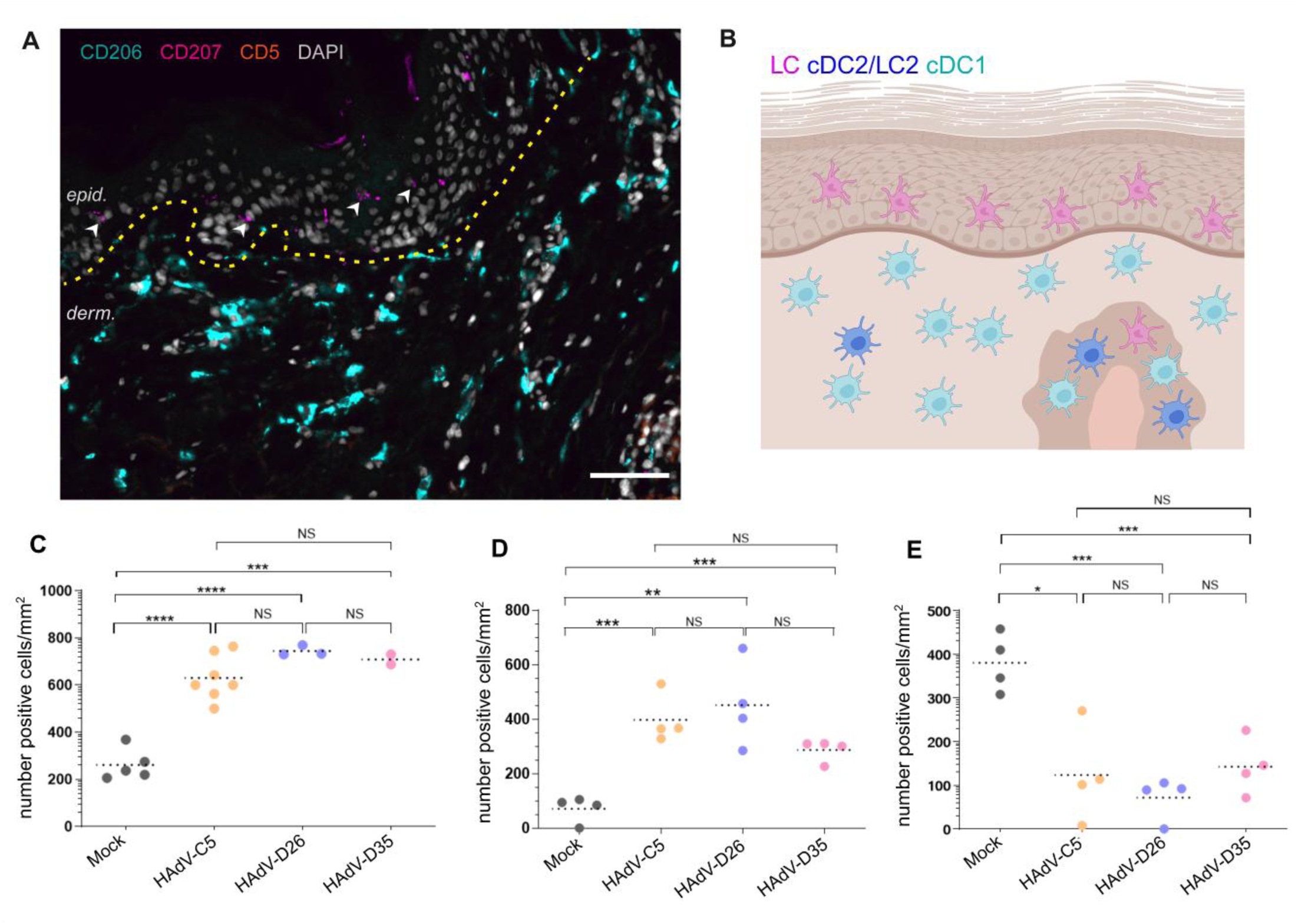
LCs and CD5^+^ APCs migrate close to the injection sites of the HAdV-based vaccine. (A) Multiplex immunofluorescence staining of human skin sections identifying resident skin DC and LCs subsets: CD207⁺ LCs (LCs), CD206⁺ CD5⁺ dermal DCs (cDC2s/LC2s), and CD206⁺CD5^-^ macrophage-like DCs (cDC1s) (scale bar = 80 μm). White arrows show LCs. (B) Schematic illustration of the distribution of the location of the LCs (magenta), cDC2/LC2 (navy blue) and cDC1 population (blue) in the different layers of human skin. (C) Quantification of CD207⁺ LCs density in the epidermis following injection of 10^11^ pp vaccine vectors. (D) Quantification of CD5⁺ cell migration toward the injection site in the epidermis. (E) Quantification of CD5⁺ cell migration toward the injection site in the dermis. The density is calculated per cell/mm^2^ using QPath software and compared to uninjected skin. A 400 µm perimeter was defined around the injection site to quantify the number of cells considered to be in close proximity to the injection area. (*p < 0.05 to ****p < 0.0001)

Given the central role of LCs in antigen uptake at the epidermal barrier and their ability to migrate to lymph nodes to initiate the immune response, we sought to determine whether injection of HAdV vectors for vaccine purposes induced local relocalization or recruitment of these cells, for subsequent antigen presentation. A dose of 10^11^ pp (physical particles) compared with the average number of pp stated in previous publications on vaccines based on HAdV-C2 and-D26 (63). Such local migration could reflect an early stage of their activation or engagement in the immune response. After injection of HAdV-C5,-D26 or-B35, a marked increase in CD207⁺ cell density was observed at the injection site (600 to 800 cells/mm² versus around 300 cells/mm² in non-injected controls), quantified using QuPath software (**Figure** 8C). This accumulation was comparable for all three vectors. Similarly, the number of CD5⁺ cells in the epidermis increased significantly after injection (**Figure** 8D), while their density in the dermis decreased (**Figure** 8E), with no notable difference between the different vectors.

These results suggest local recruitment or redistribution of CD5⁺ APCs, possibly linked to unconventional migration of cDC2s to the epidermis and/or induction of CD5 expression by primary epidermal LCs. After activation, cDC2s become LC2s, expressing langerin more weakly than the initial LC population, and thus complementing and increasing the LC population in the epidermis (64–69).

Finally, labeling with the nuclear protein Ki-67 revealed no significant difference in cell proliferation between tissues injected with HAdV vectors, PBS or healthy controls, indicating that the observed accumulation of cells does not result from local proliferation but from a migration or relocalization phenomenon (data not shown).

## DISCUSSION

LCs orchestrate the skin’s immune surveillance and form a unique population, distinguished by their ontogeny (70), function (71), phenotypic markers (e.g., langerin/CD207), and exclusive epidermal localization. As epidermal-resident APCs, these cells have long been acknowledged for their involvement in antiviral immunity. However, their contribution to vaccine-induced immune responses, particularly those caused by HAdV-based vectors, has remained poorly characterized. Here, using a dual *in vitro* and *ex vivo* approach, we demonstrate that human LCs not only detect and internalize HAdV-C5, D26, and B35 but also exhibit functional activation hallmarks that support their critical role as immune sentinels in skin-based vaccination strategies. Our results highlight the relevance of CD34-LCs as a reproducible, and physiologically relevant in vitro model for studying LC population. Despite the complexity associated with their generation, our methodology enabled consistent differentiation of over 70% CD207⁺HLA-DR⁺CD1a⁺ cells across multiple donors. Although we did not use quantitative markers such as CD80/CD86, these CD34-LCs faithfully express canonical LC markers, respond to HAdV stimuli by producing proinflammatory and antiviral cytokines, and undergo structural remodeling consistent with activation. Compared to other commonly used models moLCs, primary epidermal LCs (eLCs), or CD207-transfected CHO cells, CD34-LCs exhibit higher expression of langerin (CD207). All these elements make CD34-LCs a reliable target in the vaccination strategies as it has already been suggested with DCs (72).

We observed efficient uptake and transgene expression from all three tested HAdV vectors in CD34-LCs. The better results were especially with HAdV-B35, probably due to its affinity for certain receptors or specific MHC class I (HLA-I) allelic variants, underlining the importance of inter-individual genetic variability (73). In summary, LCs are able to capture but also secrete IFN-β, CXCL10, CCL5 and to a lesser extent IL-1β, all necessary for adaptive immunity priming and even seem to orient this response towards a Th1 profile. On the other hand, contrary to the previous results obtained with moDCs and not moLCs, transcripts of IL-6, IL-8, IFN-α, IL-18, TNF or IL-12p40 remain unchanged in the contact with HAdV-C5,-D26 and –B35, probably due to the intrinsic characteristics of LCs. Moreover, we demonstrated that HAdV exposure affects LC morphology and mechanical properties, supporting the notion that HAdV vectors trigger maturation processes in LCs.

The exploration of the Lf effect, as an immunomodulatory factor, showed that this protein can enhance HAdV uptake. Lf increased transduction efficiency of CD34-LCs across all vector types, which is particularly significant for HAdV-B35. Lf was also involved in maintaining transcript expression and cytokine secretion, particularly for CXCL10 and CCL5. Interestingly, the enhancement of LC immunodetection by Lf was not accompanied by exacerbated IL-1β secretion, suggesting that Lf does not potentiate inflammasome activation. Another important finding is that the increase in vector internalization did not induce an exaggerated immune response. The use of host-derived factors such as Lf could modulate vaccine efficacy by modifying LC-vector interactions, while avoiding a deleterious immune response. However, the alternative receptors involved in Lf-mediated internalization remains to be elucidated.

To extend these findings into a more physiological context, we used a relevant and innovant *ex vivo* model: the human skin explant, which demonstrated that epidermal LCs rapidly migrate toward HAdV injection sites and internalize vectors in situ. This initial observation reinforced our hypothesis: LCs act as first responders and enhancers in front of administered HAdV based-vaccines. First of all, we observed a redistribution of LCs into the epidermis, but also CD5⁺ APCs from the dermis into the epidermis, suggesting potential cross-talk between DC subsets. With this publication, we challenged previous representation which primarily attributed adenoviral recognition to dermal DCs and macrophages, and we placed LCs as key players in cutaneous vaccine efficacy.

In summary, our data support a new dual pathway model of adenoviral recognition by LCs: (i) a direct pathway via cell receptors like langerin, which binds HAdV capsids with serotype-specific affinities, and (ii) an Lf-mediated uptake but langerin-independent pathway.

To conclude, our study shows that LCs act in the immune detection of HAdV-based vaccine vectors in the skin. Through both direct and indirect mechanisms, LCs internalize vectors, initiate pro-inflammatory and antiviral cytokines production and adopt morphological changes, which are characteristic of immune cell activation and necessary for their ability to initiate the adaptive immune response.

The utilization of the unique properties of LCs in skin-targeted immunization strategies could lead to more effective and durable vaccine responses. In fact, LCs have recently been involved in the autonomous immune function of the skin characterized as a tertiary lymphoid organ (TLO) where they would be able to establish humoral responses independently of secondary lymphoid structures (i.e. promote T follicular helper cell differentiation and initiate the antibody responses locally) (74–76). Our research highlights the importance of LCs in the effectiveness of our vaccines targeting Skin-Associated Lymphoid Tissue (SALT) and suggest that selective targeting of Langerhans cells (LCs) by intraepidermal vaccine strategies could induce both local and systemic immune responses, and should be explored in the development of new vaccines.

We declare that we have no competing interests.

## ACKNOWLEDGEMENTS

The funders played no role in study design, data collection and analysis, the decision to publish, or preparation of the manuscript.

Study design and conception, E.G.T., FPB, EJK.; project direction, FPB & EJK; performed experiments, E.G.T, data analysis, all authors; manuscript writing, E.G.T., EJK.; securing funding, EJK.

We thank Sebastien Lyonnais from CEMIPAI, Université de Montpellier, CNRS, Montpellier, France, for AFM experiment and Thi Hong Giang Ngo from PP2I platform from IRCM for SPR analyses.

We would also like to thank Martine Pugnière from PP2I platform, Chantal Cazevieille from RHEM (INM) COMET platform, the imaging facility MRI, and RHEM histological techniques platform.

## REFERENCES

1. Sprecher E, Beeker Y. 1993. Role of Langerhans cells and other dendritic cells in viral diseases Brief Review. Arch Virol 132:1–28.

2. Kabashima K, Honda T, Ginhoux F, Egawa G. 2018. The immunological anatomy of the skin. Nature Reviews Immunology 2018 19:1 19:19–30.

3. Matejuk A. Skin Immunity 10.1007/s00005-017-0477-3.

4. Birbeck MS, Breathnach AS, Everall JD. 1961. An Electron Microscope Study of Basal Melanocytes and High-Level Clear Cells (Langerhans Cells) in Vitiligo. Journal of Investigative Dermatology 37:51–64.

5. Paul Langerhans V, reed in Berlin S. 1868. Ueber die Nerven der menschlichen Haut. Archiv für pathologische Anatomie und Physiologie und für klinische Medicin 1868 44:2 44:325–337.

6. Stingl G, Katz SI, Green I, Shevach EM. 1980. The functional role of Langerhans cells. Journal of Investigative Dermatology 74:315–318.

7. Stingl G, Tamaki K, Katz SI. 1980. Origin and Function of Epidermal Langerhans Cells. Immunol Rev 53:149–174.

8. Stingl G, Katz SI, Shevach EM, Rosenthal AS, Green I. 1978. Analogous functions of macrophages and Langerhans cells in the initiation in the immune response. J Invest Dermatol 71:59–64.

9. Kashem SW, Haniffa M, Kaplan DH. 2017. Antigen-Presenting Cells in the Skin. Annu Rev Immunol 35:469–499.

10. van den Berg LM, Ribeiro CMS, Zijlstra-Willems EM, de Witte L, Fluitsma D, Tigchelaar W, Everts V, Geijtenbeek TBH. 2014. Caveolin-1 mediated uptake via langerin restricts HIV-1 infection in human Langerhans cells. Retrovirology 11:1–9.

11. De Witte L, Nabatov A, Pion M, Fluitsma D, De Jong MAWP, De Gruijl T, Piguet V, Van Kooyk Y, Geijtenbeek TBH. 2007. Langerin is a natural barrier to HIV-1 transmission by Langerhans cells. Nat Med 13:367–371.

12. Bertram KM, Truong NR, Smith JB, Kim M, Sandgren KJ, Feng KL, Herbert JJ, Rana H, Danastas K, Miranda-Saksena M, Rhodes JW, Patrick E, Cohen RC, Lim J, Merten SL, Harman AN, Cunningham AL. 2021. Herpes Simplex Virus type 1 infects Langerhans cells and the novel epidermal dendritic cell, Epi-cDC2s, via different entry pathways. PLoS Pathog 17:e1009536.

13. Helgers LC, Keijzer NCH, van Hamme JL, Sprokholt JK, Geijtenbeek TBH. 2024. Dengue Virus Infects Human Skin Langerhans Cells through Langerin for Dissemination to Dendritic Cells. J Invest Dermatol 144:1099–1111.e3.

14. Martin MF, Maarifi G, Abiven H, Seffals M, Mouchet N, Beck C, Bodet C, Lévèque N, Arhel NJ, Blanchet FP, Simonin Y, Nisole S. 2022. Usutu Virus escapes langerin-induced restriction to productively infect human Langerhans cells, unlike West Nile virus. Emerg Microbes Infect 11:761–774.

15. Rowe WP, Huebner RJ, Gilmore LK, Parrott RH, Ward TG. 1953. Isolation of a cytopathogenic agent from human adenoids undergoing spontaneous degeneration in tissue culture. Proc Soc Exp Biol Med 84:570–573.

16. Harrach B, Benkő M. 2021. Adenoviruses (Adenoviridae). Encyclopedia of Virology 3–16.

17. Smith JG, Wiethoff CM, Stewart PL, Nemerow GR. 2010. Adenovirus. Curr Top Microbiol Immunol 343:195–224.

18. Mennechet FJD, Paris O, Ouoba AR, Salazar Arenas S, Sirima SB, Takoudjou Dzomo GR, Diarra A, Traore IT, Kania D, Eichholz K, Weaver EA, Tuaillon E, Kremer EJ. 2019. A review of 65 years of human adenovirus seroprevalence. Expert Rev Vaccines 18:597–613.

19. Fadila S, Beucher B, Dopeso-Reyes IG, Mavashov A, Brusel M, Anderson K, Ismeurt C, Goldberg EM, Ricobaraza A, Hernandez-Alcoceba R, Kremer EJ, Rubinstein M. 2023. Viral vector–mediated expression of NaV1.1, after seizure onset, reduces epilepsy in mice with Dravet syndrome. Journal of Clinical Investigation 133.

20. Li C, Psatha N, Wang H, Singh M, Samal HB, Zhang W, Ehrhardt A, Izsvák Z, Papayannopoulou T, Lieber A. 2018. Integrating HDAd5/35++ Vectors as a New Platform for HSC Gene Therapy of Hemoglobinopathies. Mol Ther Methods Clin Dev 9:142–152.

21. Schröer K, Fiege L, Wolf A, Bjelic-Radisic V, Ehrhardt A. 2025. State-of-the-art in oncolytic virotherapy using adenoviruses other than the commonly applied adenovirus type 5. Curr Opin Virol 72.

22. Ginn SL, Amaya AK, Alexander IE, Edelstein M, Abedi MR. 2018. Gene therapy clinical trials worldwide to 2017: An update. J Gene Med 20.

23. Lasaro MO, Ertl HCJ. 2009. New Insights on Adenovirus as Vaccine Vectors. Molecular Therapy 17:1333–1339.

24. Tatsis N, Ertl HCJ. 2004. Adenoviruses as vaccine vectors. Molecular Therapy 10:616–629.

25. Zhang C, Zhou D. 2016. Adenoviral vector-based strategies against infectious disease and cancer. Hum Vaccin Immunother 12:2064.

26. Chang J. 2021. Adenovirus Vectors: Excellent Tools for Vaccine Development. Immune Netw 21:1–11.

27. Zhu FC, Li YH, Guan XH, Hou LH, Wang WJ, Li JX, Wu SP, Wang B Sen, Wang Z, Wang L, Jia SY, Jiang HD, Wang L, Jiang T, Hu Y, Gou JB, Xu SB, Xu JJ, Wang XW, Wang W, Chen W. 2020. Safety, tolerability, and immunogenicity of a recombinant adenovirus type-5 vectored COVID-19 vaccine: a dose-escalation, open-label, non-randomised, first-in-human trial. Lancet 395:1845– 1854.

28. Luo H, Zhou Q, Feng J, Wu Y, Chen H, Mao M, Qi R. 2024. Global Prevalence of Preexisting Antibodies against Human Adenoviruses, Surveyed from 1962 to 2021. Intervirology 67:19–39.

29. Barouch DH, Kik S V., Weverling GJ, Dilan R, King SL, Maxfield LF, Clark S, Ng’ang’a D, Brandariz KL, Abbink P, Sinangil F, de Bruyn G, Gray GE, Roux S, Bekker LG, Dilraj A, Kibuuka H, Robb ML, Michael NL, Anzala O, Amornkul PN, Gilmour J, Hural J, Buchbinder SP, Seaman MS, Dolin R, Baden LR, Carville A, Mansfield KG, Pau MG, Goudsmit J. 2011. International seroepidemiology of adenovirus serotypes 5, 26, 35, and 48 in pediatric and adult populations. Vaccine 29:5203–5209.

30. Calabro S, Tortoli M, Baudner BC, Pacitto A, Cortese M, O’Hagan DT, De Gregorio E, Seubert A, Wack A. 2011. Vaccine adjuvants alum and MF59 induce rapid recruitment of neutrophils and monocytes that participate in antigen transport to draining lymph nodes. Vaccine 29:1812–1823.

31. Moretta A, Scieuzo C, Petrone AM, Salvia R, Manniello MD, Franco A, Lucchetti D, Vassallo A, Vogel H, Sgambato A, Falabella P. 2021. Antimicrobial Peptides: A New Hope in Biomedical and Pharmaceutical Fields. Front Cell Infect Microbiol 11:668632.

32. Chéneau C, Eichholz K, Tran TH, Tran TTP, Paris O, Henriquet C, Bajramovic JJ, Pugniere M, Kremer EJ. 2021. Lactoferrin Retargets Human Adenoviruses to TLR4 to Induce an Abortive NLRP3-Associated Pyroptotic Response in Human Phagocytes. Front Immunol 12.

33. Eichholz K, Tran TH, Chéneau C, Tran TTP, Paris O, Pugniere M, Kremer EJ. 2022. Adenovirus-α-Defensin Complexes Induce NLRP3-Associated Maturation of Human Phagocytes via Toll-Like Receptor 4 Engagement. J Virol 96.

34. Adams WC, Bond E, Havenga MJE, Holterman L, Goudsmit J, Hedestam GBK, Koup RA, Loré K. 2009. Adenovirus serotype 5 infects human dendritic cells via a coxsackievirus-adenovirus receptor-independent receptor pathway mediated by lactoferrin and DC-SIGN. Journal of General Virology 90:1600–1610.

35. Gerber-Tichet Dienst Eric Kremer EJ. 2022. Adenovirus receptors on antigen-presenting cells of the skin 10.1111/boc.202200043.

36. Keriel A, René C, Galer C, Zabner J, Kremer EJ. 2006. Canine adenovirus vectors for lung-directed gene transfer: efficacy, immune response, and duration of transgene expression using helper-dependent vectors. J Virol 80:1487–1496.

37. Malecaze F, Decha A, Serre B, Penary M, Duboue M, Berg D, Levade T, Lubsen NH, Kremer EJ, Couderc B. 2006. Prevention of posterior capsule opacification by the induction of therapeutic apoptosis of residual lens cells. Gene Ther 13:440–448.

38. Weaver EA, Barry MA. 2013. Low seroprevalent species D adenovirus vectors as influenza vaccines. PLoS One 8.

39. Tuve S, Wang H, Ware C, Liu Y, Gaggar A, Bernt K, Shayakhmetov D, Li Z, Strauss R, Stone D, Lieber A. 2006. A new group B adenovirus receptor is expressed at high levels on human stem and tumor cells. J Virol 80:12109–12120.

40. Riedl E, Stöckl J, Majdic O, Scheinecker C, Rappersberger K, Knapp W, Strobl H. 2000. Functional involvement of E-cadherin in TGF-beta 1-induced cell cluster formation of in vitro developing human Langerhans-type dendritic cells. J Immunol 165:1381–1386.

41. Tang A, Amagai M, Granger LG, Stanley JR, Uddy MC. 1993. Adhesion of epidermal Langerhans cells to keratinocytes mediated by E-cadherin. Nature 361:82–85.

42. Pranke P, Hendrikx J, Debnath G, Alespeiti G, Rubinstein P, Nardi N, Visser J. 2005. Immunophenotype of hematopoietic stem cells from placental/umbilical cord blood after culture. Braz J Med Biol Res 38:1775–1789.

43. Hamilos DL. 1989. Antigen presenting cells. Immunol Res 8:98–117.

44. Valladeau J, Ravel O, Dezutter-Dambuyant C, Moore K, Kleijmeer M, Liu Y, Duvert-Frances V, Vincent C, Schmitt D, Davoust J, Caux C, Lebecque S, Saeland S. 2000. Langerin, a novel C-type lectin specific to Langerhans cells, is an endocytic receptor that induces the formation of Birbeck granules. Immunity 12:71–81.

45. Tran TTP, Eichholz K, Amelio P, Moyer C, Nemerow GR, Perreau M, Mennechet FJD, Kremer EJ. 2018. Humoral immune response to adenovirus induce tolerogenic bystander dendritic cells that promote generation of regulatory T cells. PLoS Pathog 14:e1007127.

46. Tran TTP, Tran TH, Kremer EJ. 2021. IgG-Complexed Adenoviruses Induce Human Plasmacytoid Dendritic Cell Activation and Apoptosis. Viruses 13:1699.

47. Perreau M, Mennechet F, Serratrice N, Glasgow JN, Curiel DT, Wodrich H, Kremer EJ. 2007. Contrasting Effects of Human, Canine, and Hybrid Adenovirus Vectors on the Phenotypical and Functional Maturation of Human Dendritic Cells: Implications for Clinical Efficacy. J Virol 81:3272–3284.

48. Atasheva S, Shayakhmetov DM. 2022. Cytokine Responses to Adenovirus and Adenovirus Vectors. Viruses 14.

49. Chakraborty M, Chu K, Shrestha A, Revelo XS, Zhang X, Gold MJ, Khan S, Lee M, Huang C, Akbari M, Barrow F, Chan YT, Lei H, Kotoulas NK, Jovel J, Pastrello C, Kotlyar M, Goh C, Michelakis E, Clemente-Casares X, Ohashi PS, Engleman EG, Winer S, Jurisica I, Tsai S, Winer DA. 2021. Mechanical Stiffness Controls Dendritic Cell Metabolism and Function. Cell Rep 34.

50. Gerber-Tichet E, Blanchet FP, Majzoub K, Kremer EJ. 2025. Toll-like receptor 4 - a multifunctional virus recognition receptor. Trends Microbiol 33:34–47.

51. Eichholz K, Bru T, Tran TTP, Fernandes P, Welles H, Mennechet FJD, Manel N, Alves P, Perreau M, Kremer EJ. 2016. Immune-Complexed Adenovirus Induce AIM2-Mediated Pyroptosis in Human Dendritic Cells. PLoS Pathog 12:e1005871.

52. De La Salle H, Haegel-Kronenberger H, Bausinger H, Astier A, Cazenave JP, Fridman WH, Sautès C, Teillaud JL, Hanau D, Bieber T. 1997. Functions of Fc receptors on human dendritic Langerhans cells. Int Rev Immunol 16:187–203.

53. Junker F, Gordon J, Qureshi O. 2020. Fc Gamma Receptors and Their Role in Antigen Uptake, Presentation, and T Cell Activation. Front Immunol 11:547589.

54. Guilliams M, Bruhns P, Saeys Y, Hammad H, Lambrecht BN. 2014. The function of Fcγ receptors in dendritic cells and macrophages. Nat Rev Immunol 14:94– 108.

55. Johansson C, Jonsson M, Marttila M, Persson D, Fan X-L, Skog J, Frängsmyr L, Wadell G, Arnberg N. 2007. Adenoviruses use lactoferrin as a bridge for CAR-independent binding to and infection of epithelial cells. J Virol 81:954–963.

56. van der Aar AMG, Sylva-Steenland RMR, Bos JD, Kapsenberg ML, de Jong EC, Teunissen MBM. 2007. Loss of TLR2, TLR4, and TLR5 on Langerhans cells abolishes bacterial recognition. J Immunol 178:1986–1990.

57. Azuma H, Watanabe E, Otsuka Y, Negishi Y, Ohkura S, Shinya E, Takahashi H. 2016. Induction of langerin+ Langerhans cell-like cells expressing reduced TLR3 from CD34+ cord blood cells stimulated with GM-CSF, TGF-β1, and TNF-α. Biomed Res 37:271–281.

58. Wang H, Liaw Y-C, Stone D, Kalyuzhniy O, Amiraslanov I, Tuve S, Verlinde CLMJ, Shayakhmetov D, Stehle T, Roffler S, Lieber A. 2007. Identification of CD46 binding sites within the adenovirus serotype 35 fiber knob. J Virol 81:12785–12792.

59. Mullis KG, Haltiwanger RS, Hart GW, Marchase RB, Engler JA. 1990. Relative accessibility of N-acetylglucosamine in trimers of the adenovirus types 2 and 5 fiber proteins. J Virol 64:5317–5323.

60. Caillet-Boudin M-L, Strecker G, Michalski J-C. 1989. O-linked GlcNAc in serotype-2 adenovirus fibre. Eur J Biochem 184:205–211.

61. Cauet G, Strub JM, Leize E, Wagner E, Van Dorsselaer A, Lusky M. 2005. Identification of the glycosylation site of the adenovirus type 5 fiber protein. Biochemistry 44:5453–5460.

62. Satoh T, Otsuka A, Contassot E, French LE. 2015. The inflammasome and IL-1β: implications for the treatment of inflammatory diseases. Immunotherapy 7:243–254.

63. Afrough S, Rhodes S, Evans T, White R, Benest J. 2020. Immunologic Dose-Response to Adenovirus-Vectored Vaccines in Animals and Humans: A Systematic Review of Dose-Response Studies of Replication Incompetent Adenoviral Vaccine Vectors when Given via an Intramuscular or Subcutaneous Route. Vaccines 2020, Vol 8, Page 131 8:131.

64. Bertram KM, O’Neil TR, Vine EE, Baharlou H, Cunningham AL, Harman AN. 2023. Defining the landscape of human epidermal mononuclear phagocytes. Immunity 56:459–460.

65. Martínez-Cingolani C, Grandclaudon M, Jeanmougin M, Jouve M, Zollinger R, Soumelis V. 2014. Human blood BDCA-1 dendritic cells differentiate into Langerhans-like cells with thymic stromal lymphopoietin and TGF-β. Blood 124:2411–2420.

66. Bigley V, McGovern N, Milne P, Dickinson R, Pagan S, Cookson S, Haniffa M, Collin M. 2015. Langerin-expressing dendritic cells in human tissues are related to CD1c+ dendritic cells and distinct from Langerhans cells and CD141high XCR1+ dendritic cells. J Leukoc Biol 97:627–634.

67. Vine EE, Rhodes JW, Warner van Dijk FA, Byrne SN, Bertram KM, Cunningham AL, Harman AN. 2022. HIV transmitting mononuclear phagocytes; integrating the old and new. Mucosal Immunol 15:542–550.

68. Milne P, Bigley V, Gunawan M, Haniffa M, Collin M. 2014. CD1c+ blood dendritic cells have Langerhans cell potential. Blood 125:470.

69. Bertram KM, Botting RA, Baharlou H, Rhodes JW, Rana H, Graham JD, Patrick E, Fletcher J, Plasto TM, Truong NR, Royle C, Doyle CM, Tong O, Nasr N, Barnouti L, Kohout MP, Brooks AJ, Wines MP, Haertsch P, Lim J, Gosselink MP, Ctercteko G, Estes JD, Churchill MJ, Cameron PU, Hunter E, Haniffa MA, Cunningham AL, Harman AN. 2019. Identification of HIV transmitting CD11c+ human epidermal dendritic cells. Nat Commun 10.

70. Hoeffel G, Wang Y, Greter M, See P, Teo P, Malleret B, Leboeuf M, Low D, Oller G, Almeida F, Choy SHY, Grisotto M, Renia L, Conway SJ, Stanley ER, Chan JKY, Ng LG, Samokhvalov IM, Merad M, Ginhoux F. 2012. Adult Langerhans cells derive predominantly from embryonic fetal liver monocytes with a minor contribution of yolk sac-derived macrophages. J Exp Med 209:1167–1181.

71. Doebel T, Voisin B, Nagao K. 2017. Langerhans Cells - The Macrophage in Dendritic Cell Clothing. Trends Immunol 38:817–828.

72. Pastor Y, Ghazzaui N, Hammoudi A, Centlivre M, Cardinaud S, Levy Y. 2022. Refining the DC-targeting vaccination for preventing emerging infectious diseases. Front Immunol 13.

73. Hong SS, Karayan L, Tournier J, Curiel DT, Boulanger PA. 1997. Adenovirus type 5 fiber knob binds to MHC class I α2 domain at the surface of human epithelial and B lymphoblastoid cells. EMBO Journal 16:2294–2306.

74. Deng W, Li T. 2025. Skin as an autonomous immune organ: Antibody production and host protection. Acta Pharm Sin B 15:2795–2797.

75. Gribonika I, Band VI, Chi L, Perez-Chaparro PJ, Link VM, Ansaldo E, Oguz C, Bousbaine D, Fischbach MA, Belkaid Y. 2025. Skin autonomous antibody production regulates host–microbiota interactions. Nature 638:1043–1053.

76. Bousbaine D, Bauman KD, Chen YE, Lalgudi P V., Nguyen TTD, Swenson JM, Yu VK, Tsang E, Conlan S, Li DB, Jbara A, Zhao A, Naziripour A, Veinbachs A, Lee YE, Phung JL, Dimas A, Jain S, Meng X, Pham TPT, McLaughlin MI, Barkal LJ, Gribonika I, Van Rompay KKA, Kong HH, Segre JA, Belkaid Y, Barnes CO, Fischbach MA. 2025. Discovery and engineering of the antibody response to a prominent skin commensal. Nature 638:1054–1064.

