## Supplementary information for "Adenovirus-based vaccines transduce and activate human Langerhans cells"

### SUPPLEMENTAL MATERIAL

#### Isolation of primary epidermal LCs (eLCs) from human tissue

eLCs were isolated from human skin explants, which were manually defatted and cut into 1 $cm^2$ sections. The tissue was incubated overnight at 37°C in RPMI 1640 containing 1 mg/mL collagenase A, 20 U/mL DNase I, 1 mg/mL dispase II, 1:2,500 penicillin/streptomycin, and 2.5 µg/mL amphotericin B. The epidermis was separated and cultured in RPMI 1640 with 10% human AB serum (hAB; Sigma-Aldrich) for 48 - 72 h to collect cells in supernatant. Cells were filtered using 70 µm cell strainers, and CD1a$^+$ cells were isolated using CD1a microbeads (cat #130-051-001, Miltenyi Biotec) to obtain LCs.

#### Culture and generation of CHO-207

Chinese hamster ovary (CHO) cells were cultured in α-MEM supplemented with 10% FCS and antibiotics (100 IU/mL penicillin (cat#243.000001.02, Sigma) and 100 µg/mL streptomycin (cat#S6501, Sigma)). CHO-207 cells were generated by transfection of 0.75 µg of plasmid DNA encoding human CD207 using TurboFect transfection reagent (Thermo Fisher Scientific).

#### Isolation and differentiation of monocytes into Langerhans cells

For moLCs, we used peripheral blood mononuclear cells (PBMCs). PBMCs were isolated from healthy donors (EFS, Montpellier, France) via Ficoll-Histopaque 1077 density gradient (Sigma-Aldrich, Lyon, France) centrifugation and CD14$^+$ monocytes were isolated using magnetic separation. Finally, CD14$^+$monocytes were differentiated into moLCs with RPMI 1640 medium and the addition of 10% FCS supplemented with GM-CSF (100 ng/mL), TGF-β (10 ng/mL), and IL-4 (10 ng/mL, days 0–2; PeproTech) for 6 days.

### SUPPLEMENTAL FIGURE

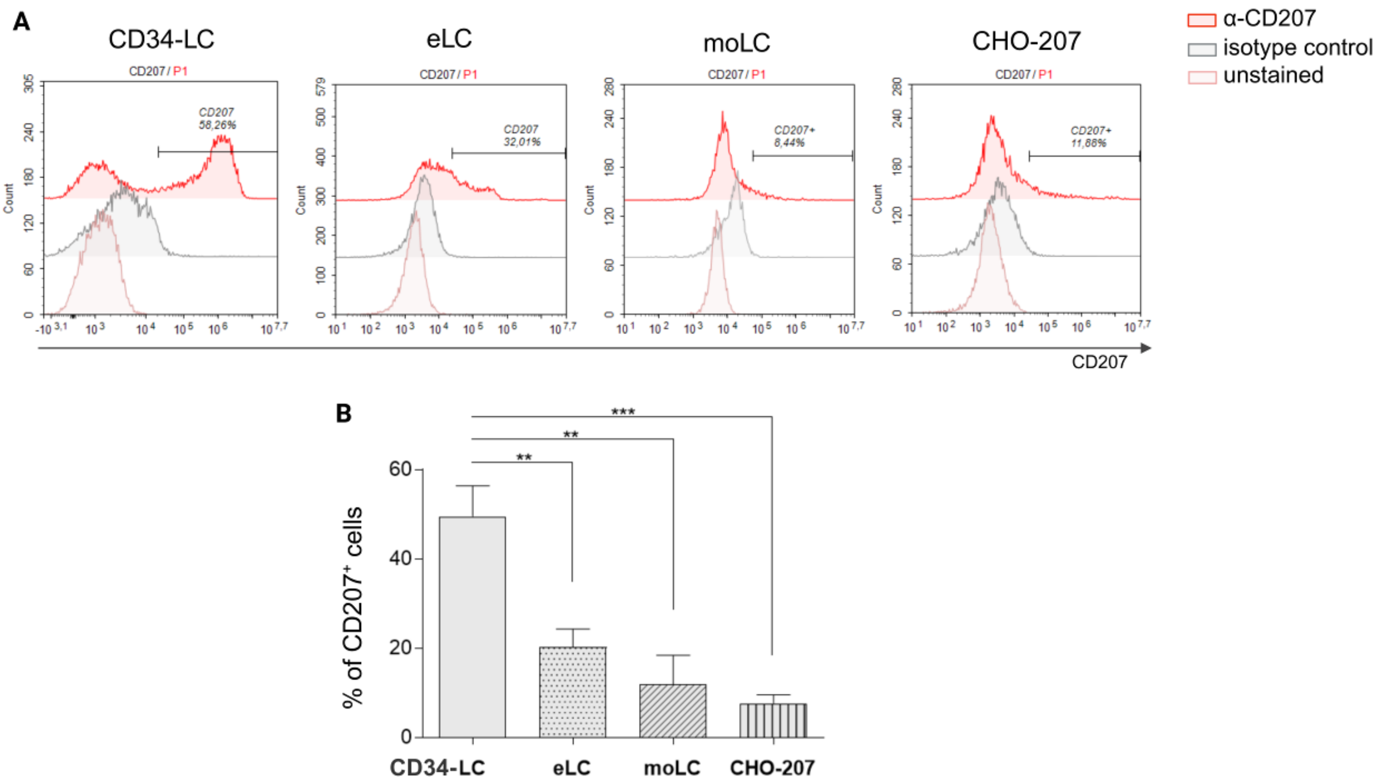


**Figure S1. LC-like models**

(A) Representative histograms showing the relative fluorescence intensity of the CD207 marker (in red) for MoDCs, moLCs, CD34-LCs, and CHO-207. Cells were stained with an anti-CD207 antibody conjugated to a fluorochrome and compared either to an isotype control (in gray) or to the condition without antibody staining (pale red). Data are representative of three independent experiments. (B) Comparative analysis of CD207 expression across different cellular models: CHO-207, moLCs, eLCs and CD34-LCs.
